# The importance of small island populations for the long-term survival of endangered large-bodied insular mammals

**DOI:** 10.1101/2024.05.23.595221

**Authors:** Sabhrina Gita Aninta, Rosie Drinkwater, Alberto Carmagnini, Nicolas J. Deere, Dwi Sendi Priyono, Noviar Andayani, Nurul L. Winarni, Jatna Supriatna, Matteo Fumagalli, Greger Larson, Peter H.A. Galbusera, Alastair Macdonald, Deborah Greer, Kusdiantoro Mohamad, Wahono Esthi Prasetyaningtyas, Abdul Haris Mustari, John Lewis Williams, Ross Barnett, Darren Shaw, Gono Semiadi, James Burton, David J. I. Seaman, Maria Voigt, Matthew Struebig, Selina Brace, Stephen Rossiter, Laurent Frantz

## Abstract

Island populations of large vertebrates have experienced higher extinction rates than mainland populations over long timescales due to demographic stochasticity, genetic drift and inbreeding. Conversely, small island populations often experience relatively less anthropogenic habitat degradation than populations on larger islands, making them potential targets for conservation interventions such as genetic rescue to improve their overall genetic diversity. Here we determine the consequences and conservation implications of long-term isolation and recent human activities on genetic diversity of island populations of two forest-dependent mammals endemic to the Wallacea archipelago: the anoa (*Bubalus* spp.) and babirusa (*Babyrousa* spp.). Using genomic analyses and habitat suitability models, we show that, compared to closely related species, populations on mainland Sulawesi exhibit low heterozygosity, high inbreeding, and a high proportion of deleterious alleles. In contrast, populations on smaller islands possess fewer deleterious mutations despite exhibiting lower heterozygosity and higher inbreeding. Site frequency spectra indicate that these patterns reflect stronger, long-term purging in smaller island populations. Our results thus suggest that conservation efforts should focus on protecting small island habitats and avoiding translocations from mainland populations. This study highlights the crucial role of small offshore islands for the long-term survival of Wallacea’s iconic and indigenous mammals in the face of development on the mainland.

**Significance statement:** Within tropical archipelagos, such as the Wallacea biodiversity hotspot, larger islands experience greater resource exploitation compared to smaller ones, highlighting the potential of smaller islands as refuges for conservation. To investigate the genetic health of populations on small islands, we used genomic, occurrence, and environmental data from a system of replicated populations of anoa and babirusa across islands of varying sizes. In contrast to larger islands like Sulawesi, our results demonstrate that smaller offshore islands not only provide higher-quality habitats but also support populations that have efficiently purged harmful mutations. Thus, despite their known vulnerability over geological time-frames, small island populations can provide long term insurance against human-driven extinction and conservation efforts should prioritise habitat management over translocations.

## Introduction

Throughout the Quaternary period, island populations of large vertebrates have experienced higher rates of extinction than their mainland counterparts^1^. This pattern can be attributed to demographic stochasticity, as well as the strong effects of genetic drift that operate in small populations of slow-reproducing species, and the associated negative consequences for fitness and adaptability^2,3^.

In the past century, these long-term threats to island populations have been further compounded by human activities such as urbanisation, the introduction of invasive species, land conversion for agriculture, mining, and hunting^4^. These anthropogenic pressures often vary across islands, with smaller and more isolated populations typically facing less intense exploitation compared to larger, more accessible ones^5,6^. Consequently, while populations on small islands may be more prone to extinction over geological timescales, they can be less affected by habitat degradation.

Among the strategies available to conservation managers of small populations, one of the most valuable is genetic rescue. In this approach, individuals from nearby, larger populations are translocated and introduced into a population to increase genetic diversity, thereby reducing inbreeding depression, mitigating the impact of recessive deleterious alleles on fitness, and potentially introducing beneficial alleles^7-9^. Although this procedure has achieved notable successes, recent empirical studies^10^ and simulations^11^ suggest that it may not always be suitable. While relocating individuals from larger populations can introduce beneficial alleles and reduce inbreeding, it also risks introducing harmful recessive alleles, potentially causing long-term negative consequences. This highlights the need for careful genetic management^12^, especially in historically small populations where natural selection has likely eliminated many harmful alleles, but which still face fitness challenges due to inbreeding and low genetic diversity.

The island archipelago of Wallacea in Indonesia provides a natural laboratory in which to determine the combined effects of long-term isolation and recent human activity on island population genetic structure and viability. Due to its long geological history of isolation, Wallacea is a global hotspot for endemism. In recent years, it has emerged as a frontier region for development in Indonesia, and its ecosystems and unique species face increasing pressures from deforestation and mineral extraction^5,6,13^. To date, human activities have been concentrated in the more accessible lowland areas on Sulawesi, whereas some smaller offshore islands have remained less disturbed.

Here we determine the consequences and conservation implications of long-term isolation and recent human activities on the genomic diversity of island populations of two forest-dependent mammals, the anoa (*Bubalus depressicornis* and *B. quarlesi*) and babirusa (*Babyrousa* spp.) (Figure 1). Both taxa that are endemic to Wallacea and are broadly co-distributed on islands of different sizes across the archipelago. Previous work suggests they underwent range expansions approximately 2 Mya^14^, however, current populations are highly fragmented following several decades of population declines and local extinctions due to habitat loss and hunting.

**Figure 1.**
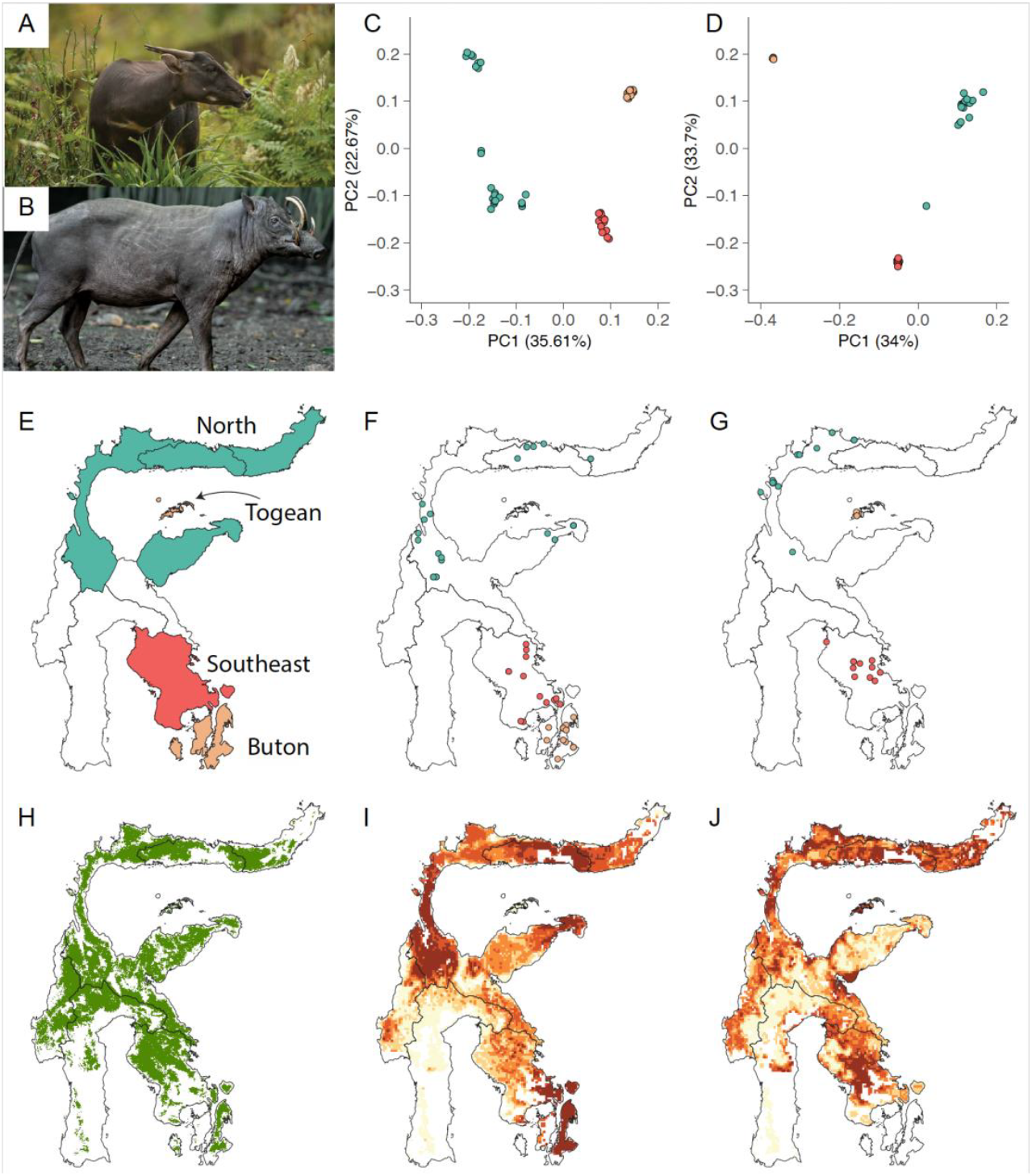
Population structure and habitat suitability. **(A)** Photographs of anoa (top, credit: Chester Zoo) and babirusa (bottom, photo credit: S.L. Mitchell). **(B)** Locations of samples for anoa (left), babirusa (right) coloured based on population structure (Figure S2). **(C)** Zones of endemism (right) based on Frantz *et al*^14^ (left) Principal component analysis (PCA) based on SNPs from genomes (>5x) for anoa (middle, 53 individuals; 1,053,534 SNPs) and babirusa (right, 37 individuals; 1,011,533 SNPs), **(D)** Sulawesi, and offshore islands, showing the 2018 forest cover (Global Forest Change repository, v1.6, Hansen *et al*^15^, processed in Voight *et al*^11^) in green used to constrain the ensemble distribution models of habitat suitability for anoa (middle) and babirusa (right) categorised by the habitat suitability score quantile (class one = least suitable habitat, class five = most suitable habitat.

As endangered and endemic mammals, the anoa and babirusa are the focus of ex-situ breeding programmes and viewed as potential candidates for conservation programmes aimed at increasing the size and genetic diversity of wild populations. Through a combination of population genomic analyses and habitat suitability models, we demonstrate that small island populations have persisted over long timeframes in high-quality habitats. In contrast, some populations on the larger island (Sulawesi; Figure 1) have suffered from reduced habitat quality due to intensive resource exploitation. Thus, conservation strategies for anoa and babirusa should focus on preserving high-quality habitats on small islands and avoid translocating individuals from degraded mainland populations. Our findings underscore the critical role of small offshore islands in ensuring the long-term survival of Wallacea’s iconic mammals amidst ongoing land-use changes on the Sulawesi mainland.

## Results and Discussion

We generated short-read genome sequence data from samples of anoa and babirusa collected from across their respective ranges in Wallacea. To determine broad population genetic structure, we mapped short reads of each taxon to a reference genome of a conspecific, or, in the case of the anoa, the water buffalo (*Bubalus bubalis*). Using datasets of 3,762,790 unlinked SNPs for anoa, and 3,920,000 unlinked SNPs for babirusa, we performed principal component analysis (Figure 1C), phylogenetic reconstructions, and admixture analyses (Figure S2). These analyses revealed concordant patterns of population structure across both taxa (Figure 1C), in which three distinct lineages were identified corresponding to individuals from North Sulawesi, Southeast Sulawesi, and the small offshore islands of Buton (anoa) or Togean (babirusa). This pattern of population structure, which broadly supports previous findings based on mitochondrial and microsatellite data^14^, was then used to define populations in subsequent analyses.

### Populations on smaller island are less genetically diverse than on mainland Sulawesi

To determine the consequences of long-term isolation, we quantified levels of genetic diversity in populations on Sulawesi and smaller nearby islands (Buton and Togean). To do so, we computed genome-wide Watterson’s *θ* using ROHan^16^, an unbiased estimator of heterozygosity in a single diploid individual under the infinite sites model^17^. We found that *θ* on the small islands of Togean (babirusa: mean=0.29 × 10^−3^, SE=0.02 × 10^−3^) and Buton (anoa: mean=0.58 × 10^−3^, SE=0.02 × 10^−3^) was ~2 to ~6 fold lower than in populations from mainland Sulawesi (babirusa: mean=1.85 × 10^−3^, SE=0.1 × 10^−3^; anoa: mean=1.33 × 10^−3^, SE=0.1 × 10^−3^), indicating that small island populations contain less genetic diversity than populations on the larger island of Sulawesi (Figure 2A). On Sulawesi, however, we found that babirusa from the Southeast Sulawesi possessed lower *θ* (mean=1.17 × 10^−3^, SE=0.04 × 10^−3^) compared to those found on the Northern peninsula (mean=2.17 × 10^−3^, SE=0.06 × 10^−3^).

**Figure 2.**
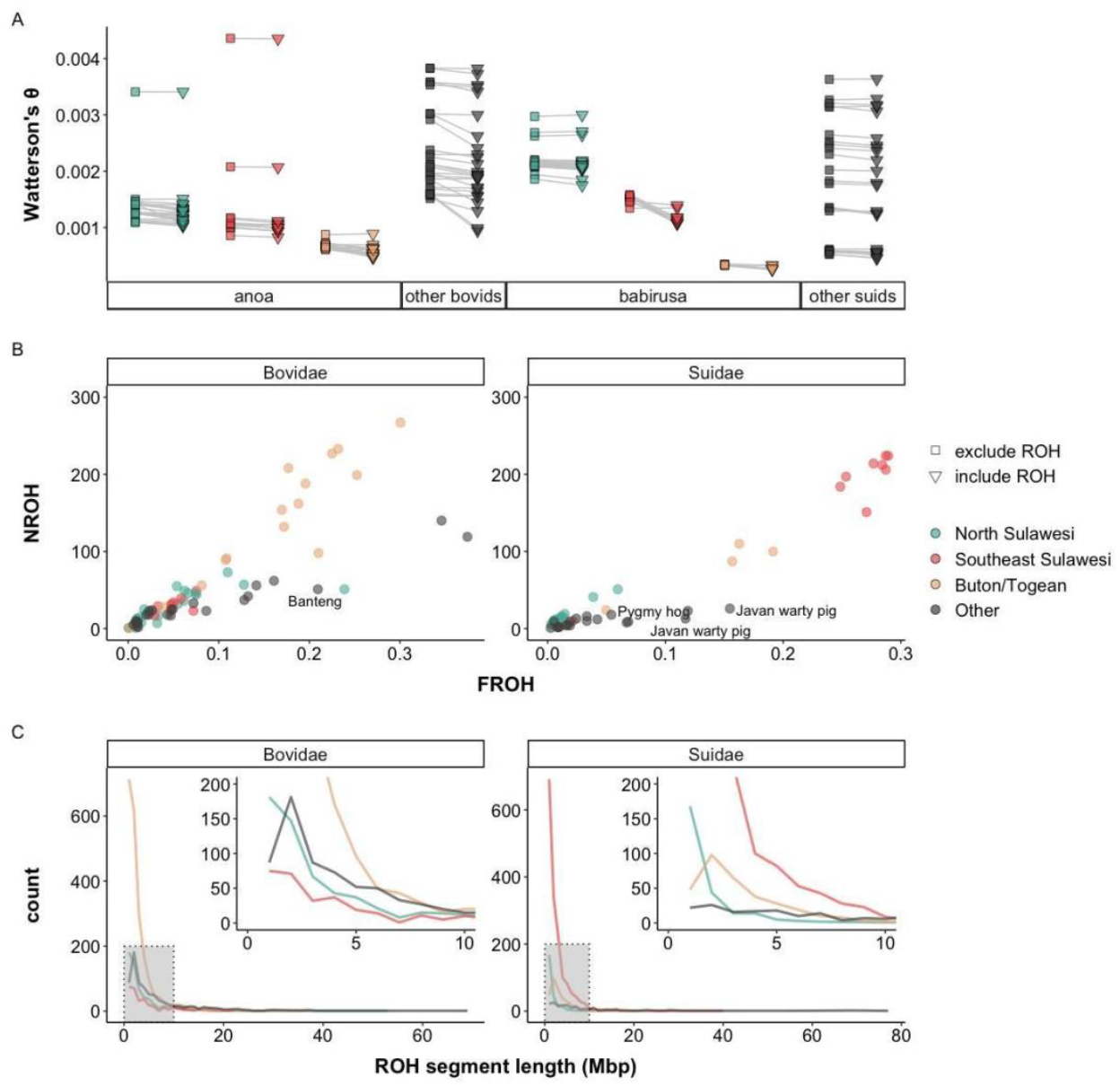
Population-level genetic diversity and runs of homozygosity (ROHs). (**A**) Genome-wide Watterson’s *θ* computed across the whole genome (triangle) and excluding ROH (square) in anoa, babirusa, and their respective close relatives, **(B)** Sum of size of ROH segments across the genome (NROH) and proportion of ROH across the genome (FROH) **(C)** distribution of ROH segment length.

We then tested whether anoa and babirusa populations are characterised by more variable levels of genetic diversity compared to related species that do not occur on islands of varying size. To address this, we compared levels of genetic variability (Watterson’s *θ*) and inbreeding (ROHs) among anoa and babirusa populations to those observed in 19 closely related taxa (52 individuals in total), including highly endangered species such as the pygmy hog (*Porcula salvianus*), the Javan warty pig (*Sus verrucosus*) and banteng (*Bos javanicus*). Levels of genetic diversity (Watterson’s *θ*) were on average lower in anoa than in other bovid species (Figure 2A). The same was observed in babirusa, which aside from the North Sulawesi population, possessed lower levels of genetic diversity than most other suids. Lower levels of genetic diversity in the Wallacean endemics is likely the result of smaller effective population sizes resulting from their island habitats and/or recent bottleneck(s).

To assess whether populations showed signs of recent bottlenecks, we analysed runs of homozygosity (ROHs) using ROHan (Supplementary Information). Analyses of ROHs showed that inbreeding levels and genetic diversity were more variable among anoa and babirusa populations than in different species (Figure 2B). Small island populations of anoa (Buton) and babirusa (Togean) possessed higher levels of inbreeding than most other species (Figure 2B). Buton contained longer ROHs than those sampled on mainland Sulawesi, consistent with higher inbreeding levels on small offshore islands (Figure 2C).

Among suids, however, we found that Southeast Sulawesi babirusa possessed more ROHs and had a larger portion of their genome in ROH than any other population of babirusa and any other suid species (Figure 2B). Interestingly, despite this high level of inbreeding, Southeast Sulawesi babirusa possessed average levels of genetic diversity when compared to other suid species (Figure 2A). Their level of genetic diversity, however, was significantly higher when excluding ROHs (p<0.001; Figure 2B). The significant change in genetic diversity when excluding ROHs implies that babirusa living in Southeast Sulawesi were part of a historically large population which underwent a recent bottleneck.

Similar levels of genetic diversity within and outside ROHs in anoa and babirusa living on small islands (Togean and Buton) indicate they form part of a historically small, yet stable population that inhabits a highly suitable habitat, as demonstrated by our species distribution model (Figure 1D, Figure S4, Table S2). Altogether, this suggests that although they are not as genetically diverse as their large island counterparts, small island populations are likely to have been stable over geological time scales, and potentially since their expansion ~2Mya^14^.

### Impact of anthropogenic disturbances on demography

To address whether the dramatic bottleneck in Southeast Sulawesi babirusa is consistent with a recent decrease of suitable habitat in the area, we constructed an ensemble species distribution model^18^ based on bioclimatic variables and forest cover^5,15^ (Supplementary Information). Our models, for both anoa and babirusa, indicate that there is proportionally more suitable habitat (i.e. top 20% of all suitable habitat; see Supplementary Text) on smaller islands (i.e. Togean and Buton) compared to mainland Sulawesi (~1.5 fold more highly suitable area for anoa and ~1.6 for babirusa relative to North Sulawesi; Figure 1D, Table S3).

For both species, we related the distribution of suitable habitat to different land use classes and found considerable overlap with areas that have protected status (Figure S4). This highlights the effectiveness of the protected area network for conserving habitat for Wallacea’s endemic mammals. On the mainland, the availability of suitable habitat was higher in the North compared to the Southeast for both taxa (Table S3). This trend is consistent with recent deforestation rates (between 2000-2017), which were higher in Southeast Sulawesi compared to the North^19^, suggesting that deforestation has led to the reduced habitat availability detected by our models. These data, combined with previously reported high rates of poaching^20^, imply that population bottlenecks in babirusa in Southeast Sulawesi have been driven by relatively recent anthropogenic disturbances, rather than longer term evolutionary processes.

### More deleterious alleles segregate on Sulawesi than on small islands

Recent studies of several large-bodied mammals indicate that small populations with long histories of isolation (*e*.*g*. Channel Island foxes^21^, Iberian lynx^22^, Indian tigers^23^, mammoths^24^) show evidence of accumulation of mildly deleterious alleles, while naturally purging strongly deleterious recessive variants. To evaluate how small island populations of anoa and babirusa purge deleterious alleles compared to their mainland populations, we computed genetic load in individual genomes, using three conservation scores: SIFT, PhyloP and phastCons (Supplementary Information). To allow cross-species comparisons, we estimated genetic load for loci that show one-to-one orthology between the pig and cow genomes (Supplementary Information). We first computed a load score by summing conservation scores, weighted by genotype probability for derived alleles found at homozygous states across the genome (see Supplementary Methods). This homozygous load represents the minimum impact of deleterious alleles on fitness (i.e. assumes that all deleterious alleles are recessive). We also computed the load by summing the impact of alleles found at both homozygous and heterozygous states (total load) across the genome. The total load represents the maximum impact of deleterious alleles on fitness (i.e. assumes that all deleterious alleles are dominant).

Both total and homozygous load were significantly lower in anoa than in babirusa (p<0.001; see Supplementary Materials) (Figure 3A & B). Mean total load, calculated for all individuals from the same population, was also significantly higher in large populations found on mainland Sulawesi than in smaller island populations such as Togean (babirusa) and Buton (anoa; Figure 3C, Table S1). This result indicates that deleterious alleles are more abundant in larger, mainland, populations than in smaller, island populations.

**Figure 3.**
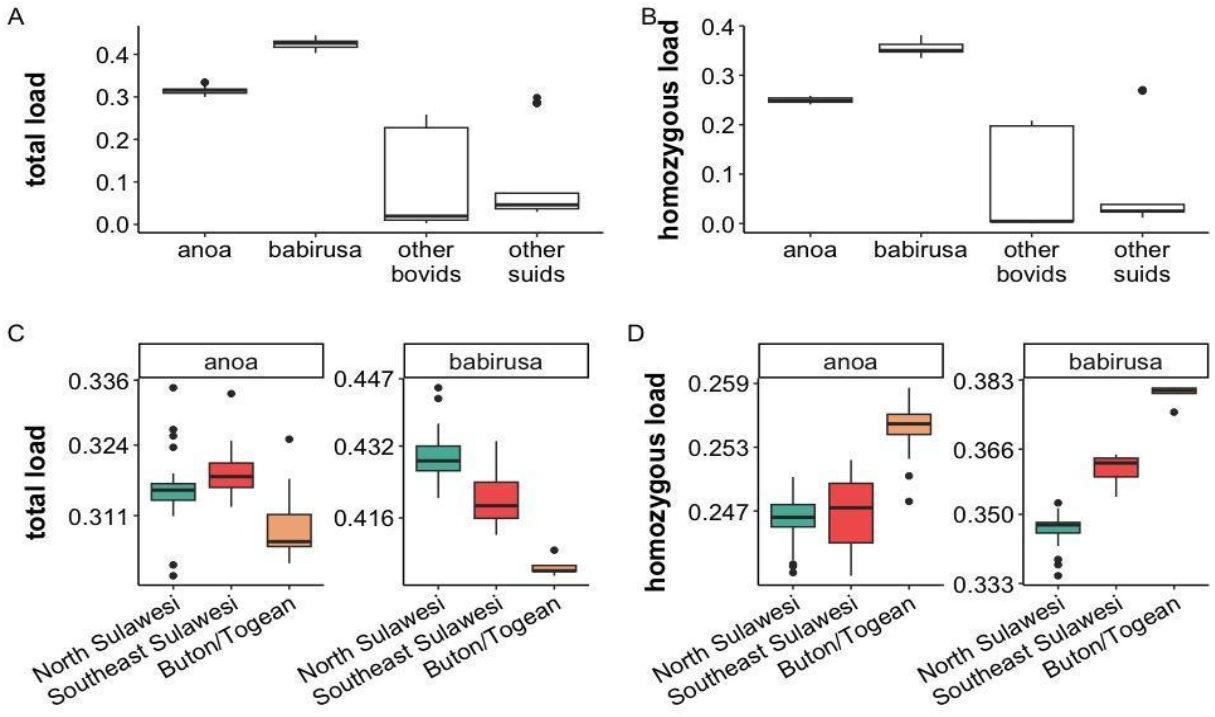
Genetic load in babirusa, anoa and closely related species. **(A)** Total load computed in individual genomes using SIFT scores for alleles found at both heterozygous and homozygous states in anoa, babirusa, and their close relatives. (**B**) Homozygous load computed in individual genomes using SIFT scores for alleles found at homozygous states in anoa, babirusa, and their close relatives. **(C)** Inter-island comparison of total load and **(D)** homozygous load within the populations of anoa and babirusa.

Mean homozygous load, however, was significantly higher in small island populations than in larger island populations (Figure 3D). Altogether these results indicate that large populations possess more deleterious recessive alleles overall, most of which are found in heterozygous state and therefore hidden from selection. Thus, although large populations possess more deleterious alleles, the overall fitness impact of deleterious alleles is likely to be higher in smaller island populations which possess more deleterious alleles in homozygous state.

### Small island populations efficiently purge deleterious alleles

The difference between the total and homozygous load observed in small and large populations could be due to either lower levels of heterozygosity, or the lower efficiency of purifying selection in small populations (Figure 2B). To assess the efficiency of purifying selection across different populations, we first built unfolded site frequency spectra (SFS) using ANGSD^25^, based on alleles that have been assigned different impact ratings by the Variant Effect Predictor (VEP)^26^, *i*.*e*. low, modifier, moderate, and high (Figure 4, Supplementary Information). Comparing SFS provides the opportunity to assess the impact of purifying selection on the full frequency range of deleterious alleles in a population, not just those in heterozygous and homozygous states.

**Figure 4.**
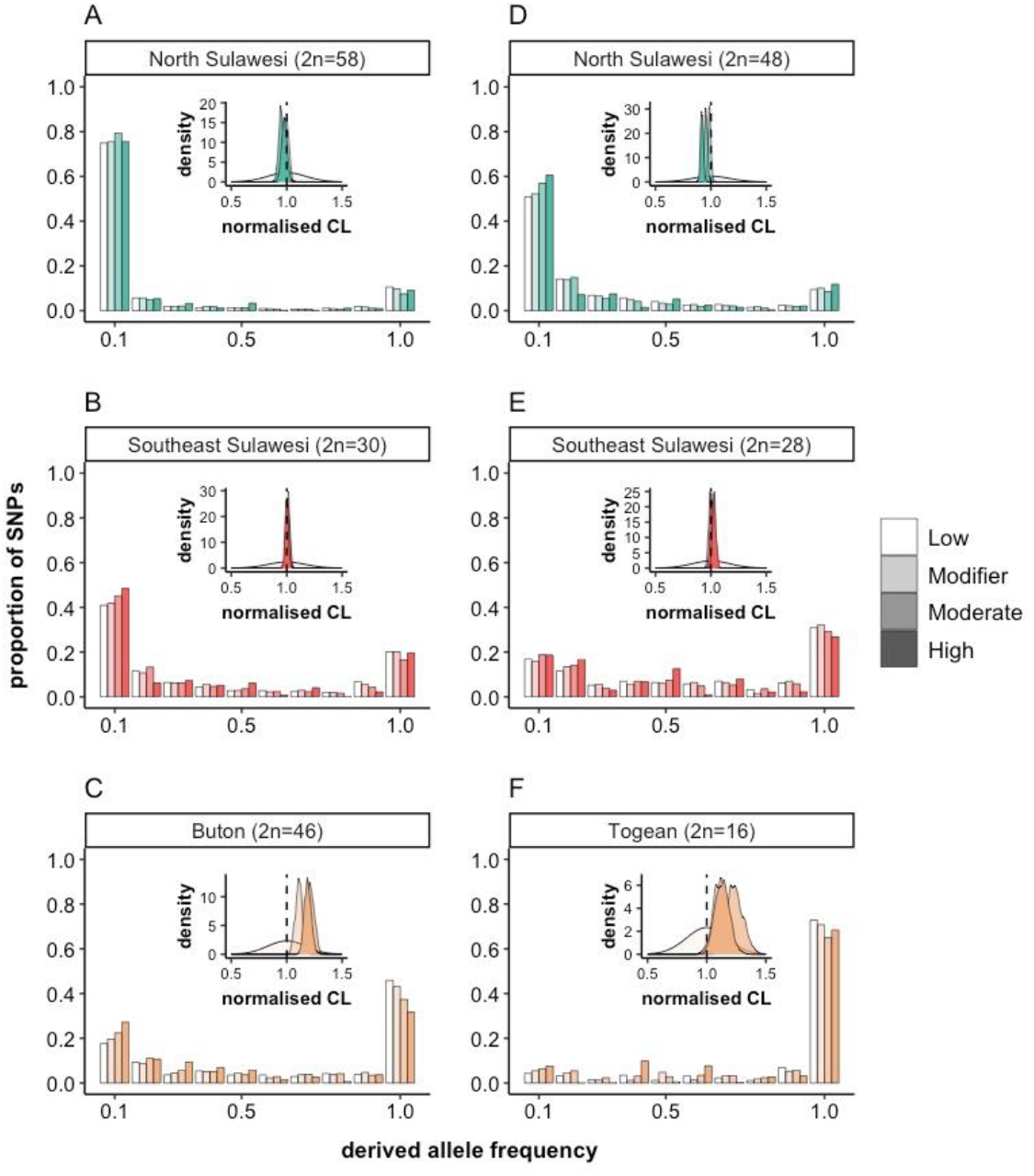
Site frequency spectra (SFS) of deleterious alleles across anoa and babirusa populations. Site frequency spectra of alleles with four different impact ratings (low, modifier, moderate, high) across populations of anoa **(A-C)** and babirusa **(D-F)** with a corresponding 1000 bootstrap of composite likelihood (CL) values within each SFS graph representing differences between the shape of the SFS across the four impact ratings using low impact rating as the expected likelihood.

These SFS show an excess of low frequency deleterious alleles in mainland populations that have not experienced bottleneck(s), such as populations of anoa in Southeast Sulawesi (Figure 4A & B) and of both taxa in North Sulawesi. This pattern is consistent with the effect of purging reducing the frequency of deleterious alleles in a population^21-23^. The persistence of deleterious alleles at low frequency in these large, highly heterozygous, populations is likely due to their recessive nature which means they are less likely to be exposed to selection than in smaller, less heterozygous, populations

In contrast, we found fewer deleterious alleles at low frequency in the recently bottlenecked Southeast Sulawesi babirusa, compared to in the North Sulawesi population. This is likely due to a weaker effect of purging, and stronger drift in Southeast Sulawesi babirusa than in the North Sulawesi. Both small island populations, however, possessed many fixed deleterious alleles, but had fewer deleterious alleles at low frequency than in large populations and in the recently bottlenecked Southeast babirusa population. The high degree of fixed deleterious alleles is likely to be the result of strong genetic drift in small populations. Reduced heterozygosity in these island populations likely explains the near absence of low-frequency deleterious alleles, a result of long-term purging where recessive alleles are exposed to selection. This is in contrast with large populations, in which deleterious alleles persist at low frequency despite purging.

To quantitatively compare the shape of the SFS across four impact ratings, we used a composite likelihood (CL) approach adapted from Nielsen *et al*^27^. Large differences in CL values across impact ratings point to contrasting shapes of the SFS. As the low impact alleles are less likely to be affected by selection compared to the other three impact ratings (modifier, moderate, high), we used the SNPs in this rating as the expected SFS. For each impact rating we obtained 1000 CL values across bootstrap replicates (see supplementary methods), which were normalised using the mean of the bootstrap value of the low impact CL. In large island populations, normalised CL distributions (across bootstrap replicates) of the three higher impact ratings were centred around one (Figure 4A, B, D, E). This indicates that the SFS of the higher impact ratings were not quantitatively different from that of the SFS built using low impact ratings. In contrast, in small island populations, the mean CL across the three higher impact ratings was higher than one, indicating that the shape of the SFS of the higher impact alleles are quantitatively different from the shape of the SFS built using low impact alleles (Figure 4C & D).

These differences in the shape of the SFS across different impact ratings in the small island populations suggests that although they are smaller than their mainland counterparts, purging is having a stronger distorting effect on the SFS of more impactful alleles. This could be the result of recessive deleterious alleles that are more likely to be exposed to selection in a less heterozygous population. Over time, this will lead to a reduction in frequency for the most deleterious alleles in small populations, consistent with our total load scores (Figure 3A) and previous simulation studies^11^.

## Conclusions

Our analyses show that anoa and babirusa populations on small islands have remained sufficiently stable to efficiently purge deleterious alleles. These populations also occupy high-quality habitats, often within protected areas (e.g. 45% of the Togean islands are protected as a national park). Small island ecosystems may thus offer a long-term solution to preserve these species. In contrast, we show that some populations on the larger island of Sulawesi, such as in the Southeast, occupy lower quality habitat due to a higher degree of anthropogenic disturbance, and show signs of strong bottlenecks and weaker purifying selection. These findings imply that recent anthropogenic disturbances on larger islands may be reshaping the extinction dynamics during the Quaternary, a period during which populations on smaller islands have been generally presumed to be more vulnerable. Our results suggest a reversal of this trend, which when combined with a greater human impact could make large island populations more susceptible to extinction than their smaller island counterparts.

The introduction of individuals from the mainland could be a solution to increase genetic diversity on small islands with high habitat quality and low anthropogenic pressures. Our genomic data, however, indicate that individuals from mainland populations possess more deleterious alleles, which, if translocated to smaller, less heterozygous, island populations could result in fitness decline and increased risk of extinction^10^. We therefore suggest that, unless the population size of babirusa and anoa drops dramatically on smaller islands, conservation efforts should focus on maintaining forest habitat, without the need for a possibly counterproductive, and onerous, translocation programme. Translocations, however, could also become useful in this case if the degree of fixed deleterious alleles, which is higher in smaller island populations (Figure 4), starts having a strong impact on fitness, as purifying cannot remove these alleles once they are fixed, or to enhance immune function by introducing new alleles. Altogether, our study demonstrates the benefit of combining genomic information with species distribution modelling to help predict future anthropogenic threats and inform species conservation planning for island systems.

## Materials and Methods

Samples used for this study were obtained from a previous study^14^ totalling to 67 anoa (29 from North Sulawesi, 15 from Southeast Sulawesi, and 23 from Buton) and 46 babirusa (24 from North Sulawesi, 14 from Southeast Sulawesi, and 8 from Togean). Detailed descriptions of the samples used for this study are presented in SI Appendix. DNA was extracted either from hair follicles from tails or tissue scraps from skeletal remains using the DNeasy Blood and Tissue kit (Qiagen) with the final extract eluted in 100 μL of TE buffer. Double indexed standard illumina libraries were built by Novogene (in 2020) or Macrogen (in 2021) (Data S1). Libraries were pooled equimolarly and sequenced on a Illumina Novaseq S4 platform (150 bp PE). To compare with other taxa that are closely related, we downloaded 25 genomes from European Nucleotide Archive (ENA) and 26 genomes from Sequence Read Archive (SRA) comprising 8 species of Suidae and 10 species of Bovidae (Data S2) that were also sequenced using Illumina (150 bp PE). Each paired end fastq files sample was trimmed with AdapterRemoval^28^ and aligned to the using the BWA MEM^29^ to a closely related reference genome, i.e. water buffalo and babirusa, and distantly related reference genome, i.e. cow and pig, for anoa and babirusa, respectively, constructing each to a set of close relative alignment and distant relative alignment (see SI Text). Other than the population structure and genetic load analyses that use the distant relative alignment to get gene annotation information, all downstream analyses were conducted using the close relative alignment. Only genomes with mean reads depth of at least 5x were analysed. Habitat suitability models were generated using anoa and babirusa occurrence data and environmental covariates (see SI text).

Ensemble distribution models^30^ were generated and analysed in R^31^ and QGIS^32^. Further details for computational analyses used in this study are provided in SI Appendix.

## Acknowledgement

This research was funded by the NERC-Ristekdikti Newton Wallacea joint research programme (grant number NE/S007067/1; Ristekdikti Grant No: NKB-1799/UN2.R3.1/HKP.05.00/2019 “Biodiversity, Environmental Change, and Land-use Policy in Sulawesi and Maluku”). We thank the Indonesian Ministry of Environment and Forestry (KLHK), Sulawesi’s Provincial Forestry Departments (BKSDA), and Indonesia’s National Research and Innovation Agency (BRIN, formerly LIPI) for granting research and export permits. This work is a continuation of research under the permit from Ministry of Forestry no 188/Kpts-IV/2001 and 2918/Kpts-IV/2002, now extended under permit no.

SK.18/KSDAE/SET-3/KSA-2/I/2024. This research was supported by computational and data resources provided by the Leibniz Supercomputing Centre (LRZ; www.lrz.de). We thank Athena Syarifa for her support during the start of the project, and Gabriel Renaud for assistance in ROH analysis. SGA was supported by QMUL and GCRF. RD is supported by NERC and Humboldt Foundation. MV, NJD and MJS were supported by a Leverhulme Research Leadership Grant Awarded to MJS. SB, SR, and LF were supported by NERC Newton Fund Wallacea. JB was supported by University of Edinburgh, Rufford Grant, and Stichting Dierentuin Helpen Grant. PG acknowledges the structural support of the Flemish government.

## Data sharing

All sequences used in this study are stored in European Nucleotide Archive accession ERS10527733-ERS10527854. All codes used for analyses in this paper is openly available in https://github.com/sagitaninta/Wallacea (see Supplementary Information)

