## Supplementary Information for "The importance of small island populations for the long-term survival of endangered large-bodied insular mammals"

**This PDF file includes:**

Supporting text
Figures S1 to S5
Tables S1 to S4
Legends for Datasets S1 to S3
SI References

**Other supporting materials for this manuscript include the following:**

Datasets S1 to S3

### Supporting Information Text

#### Methods-Sequencing Data Preparation

**DNA extraction and sequencing.** The 67 anoa and 46 babirusa samples resequenced in this study were initially collected in early 2000s[1-3] (Data S1). For anoa, a total of 25 samples originated from North Sulawesi (classified as North Central=3, Northwest=9, West Central=13, East Central=4, in Frantz *et al.*[3]), 15 from Southeast Sulawesi, and 23 from Buton. For babirusa, a total of 24 samples came from North Sulawesi (classified as Northwest=11, West Central=13 in Frantz *et al.*[3]), 14 from Southeast Sulawesi, and 8 from Togean.

DNA was extracted either from hair follicles from tails or tissue scraps from skeletal remains using the DNeasy Blood and Tissue kit (Qiagen) with the final extract eluted in 100 µL of TE buffer. Anoa DNA extractions were conducted at the University of Edinburgh (by James Burton) between 2001-2004, and the babirusa DNA extractions were conducted in the Faculty of Veterinary Medicine, Bogor Agricultural University (IPB), Indonesia (by Kusdiantoro Mohamad and Wahono Esthi Prasetyaningtyas), and in the Royal Zoological Society of Antwerp (RZSA), Belgium (by Peter Galbusera), between 2005-2006. Double indexed standard illumina libraries were built by Novogene (in 2020) or Macrogen (in 2021) (Data S1).

Libraries were pooled equimolarly and sequenced on an Illumina Novaseq S4 platform (150 bp PE). This resulted in between 9.7 and 255.9 million read pairs from 69 anoa libraries (there are two samples with two corresponding libraries) and between 16.2 and 381.4 million read pairs from 47 babirusa libraries (one sample with two libraries) (Data S1).

**Read processing and alignment.** Paired reads were trimmed for adapters and quality using AdapterRemoval v2.3.1[4] and aligned with bwa-mem v0.7.17-r1188[5]. For each taxa, we aligned the reads to two sets of reference genomes using BWA mem with default settings. First, we mapped anoa reads to the cattle reference genome (*Bos taurus* ARS-UCD1.2, also known as Btau9) and babirusa reads to the pig reference genome (*Sus scrofa*, Sscrofa11.1, also known as Sscr11). The reference genomes were downloaded from Ensembl[6]. This alignment is referred to as the "distant relative alignment" throughout the manuscript.

We then mapped the anoa reads to a chromosome level assembly of the water buffalo (*Bubalus bubalis*, UOA_WB_1, GCF_003121395.1) downloaded from Ensembl[6] Rapid Release and the babirusa reads to a draft, chromosomal length, assembly of a Sulawesi babirusa (*Babyrousa celebensis*). The babirusa reference genome was built from short insert-size PCR-free DNA-Seq data using w2rap-contigger[7] by the DNA Zoo team (see Dudchenko *et al.*[8] for more details). This chromosome-length assembly is available on DNA Zoo portal[9]. This alignment is called the "close relative alignment".

In both alignments, duplicate reads were removed using Picard MarkDuplicates[10]. To assess the quality of the two alignments, we calculated the mean reads mapped and breadth of coverage using 'samtools depth'[11]. We then compared these summary statistics on both distant and close relative alignments to assess whether the use of different reference genomes impacted the quality of the local alignment.

Percentage of reads mapped to the close relative reference genome were generally higher in both anoa and babirusa genome compared to mapping to distant relative reference genome (see Dataset S1). On average, alignments to close relatives resulted in higher mean read depth (6.9% higher for anoa and 0.7% higher for babirusa on average) and higher breadth of coverage (4.16% higher for anoa and 2.23% higher for babirusa (Figure S1, also see Dataset S1 for values of each sample). Mapping to a close relative reference genome increased the breadth of coverage in 111 out of 113 samples (65 anoa and 46 babirusa, see Dataset S1).

We used the alignments to the close relative for all downstream analyses except for the phylogenetic, population structure and the genomic load analyses to make use of the better annotation of the cow and pig genome. For the close relative alignment, the resulting coverage ranged between ~0.04x to ~21x for anoa and ~0.2x to ~29x for babirusa, and 54 anoa and 36 babirusa samples had a coverage of at least 5x (Dataset S1). All samples were used for all of the downstream analyses except the population structure analyses, ROH, and genomic load calculation with SIFT scores where only genomes with at least a 5x mean read depth were used.

**Publicly available data.** We downloaded 25 genomes from European Nucleotide Archive (ENA) and 26 genomes from Sequence Read Archive (SRA) comprising 8 species of Suidae and 10 species of Bovidae (Dataset S2). The resulting fastq reads were run through the same pipeline as anoa and babirusa, excluding the adapter trimming step.

**Methods – Downstream Analyses**

**Population structure analyses.** Samples with 5x greater coverage based on distant relative reference alignment (53 anoa and 37 babirusa, Dataset S1), and an outgroup for each taxon (anoa = african buffalo, SRA accession: SRR12191781, babirusa = Pygmy hog, ENA accession: ERR2984766) were used for the SNP analyses. Variants were called across autosomal chromosomes using graphtyper[12] using alignments to the distant relative each for anoa and babirusa. VCF were merged and indexed using bcftools[11]. We kept only biallelic SNPs flagged as "FILTER=PASS", with a genotype quality of 30, average sequencing depth of four and allowing a 10% missing genotype threshold, using bcftools. VCFs were converted into Plink format, version v1.90b6.2[13], and pruned for minimum allele frequency (*--maf 0.05*), linkage disequilibrium (*--indep-pairwise 50 5 0.5*) and thinned (*--bp-space 1500*) aiming for approximately 1 million SNPs. This resulted in 1,053,534 SNPs for anoa and 1,011,533 SNPs for babirusa that were used for ancestry analyses. We used EMU[14], specifying four eigenvectors to generate PCAs (removing outgroup species (Figure 1C).

The filtered plink .bed file was used to calculate admixture for both taxa using the software ADMIXTURE[15]. Six values of K were tested (K2-7) and 10 replicates were conducted for each cluster value. Cross-validation error was calculated with the --cv- flag, where a lower cv error value indicates that the K value is better fitting for the data. The ten replicates for each K were merged using the '*mergeQ( )*' function and plotted using POPHELPER[15] in RStudio[16] with R version 4.3.318[17].

Filtered VCFs were converted into a phylip format using vcf2phylip v2.0[19]. IQ-TREE version 2.0.3[20] was used to build phylogenies for each taxon. First invariant sites were removed, with IQ-TREE and the best fitting substitution model was found using ModelFinder[21], restricted to search only ascertainment bias correction models (-m MFP+ASC). Trees were then run using the best fitting model based on BIC (anoa = GTR+F+ASC+R2, babirusa = TVM+F+ASC+R2) and support was assessed using 1000 ultrafast bootstraps replicates[22], and 1000 replicates of the single branch test, SH-like approximate likelihood ratio[23]. The trees (Figure S2) were then plotted using the *plot.phylo.to.map* function from the "phytools" package[24] in RStudio[17] with R version 4.3.3[18].

**Identifying Runs of Homozygosity (ROH).** We used ROHan[25] to identify Runs of Homozygosity (ROH) in autosomal chromosomes. The number of segregating sites allowed in ROH (--rohmu) was set to $4x10^{-5}$ (i.e. Ne of ~1000 and $1x10^{-8}$ mutation rate) and the minimum sliding window is 1 Mbp following the default. It is also to avoid short segments that are more likely to be false positives coming from extra variants commonly appearing from mapping to different species that are breaking down the actual ROH[26]. ROHan was run directly on BAMs from the close relative alignment on genomes with a mean reads depth of at least 5x following the software recommendation. Mappability filter for each reference genome was built following the SNPable pipeline[27]. Briefly, we split the reference genome fasta into fragments of 500 bp and aligned these fragments to the same reference genome using BWA aln'[27] with the default penalty parameters to filter out unmappable regions specified by Ceballos *et al*.[28], i.e. `-R 1000000 -O 3 -E 3`. A BED file containing the coordinates of mappable regions was then provided to ROHan using the `--map` option.

We used midpoint estimates to calculate mean length of ROH segments (LROH), the genomic inbreeding coefficient (FROH), and the number of ROH segments (NROH) to summarise the ROH distribution within each individual[28]. LROH was computed per individual as:

$$L_{ROH} = \frac{\sum_{k=1}^{n_{ROH}} \quad l_{ROH_k}}{N_{ROH}} \tag{1}$$

where $N_{ROH}$ is the total count of ROH segments for in an individual and $l_{ROH_k}$ is the length of each $k^{th}$ ROH (in base pairs). The inbreeding coefficient (FROH) for each individual was then calculated as:

$$F_{ROH} = \frac{\sum_{k=1}^{n_{ROH}} \quad l_{ROH_k}}{l_g} \tag{2}$$

where $l_g$ is the total length of mappable regions in the (autosomal) genome. The minimum cutoff to calculate FROH follows the shortest ROH we detected using ROHan, which is 1Mbp. We found that the FROH becomes much smaller for longer segment groups only in inbred populations such as Southeast Sulawesi and Togean for babirusa and Buton for anoa, while outbred populations such as North Sulawesi for both and Southeast Sulawesi for anoa did not have change in FROH with increasing segment length (Fig S3, Table S1).

**Polarising alleles.** We polarised alleles for genomic load analysis (i.e. infer derived or ancestral state at each SNP). To do so, we aligned the genome assembly of three outgroups for each Bovidae (anoa and its close relatives) and Suidae (babirusa and its close relatives) to the reference genomes used for short reads. For Bovidae outgroup we included the following taxa:

| | Species name | NCBI Genome Accession |
|---|---|---|
| Reindeer | *Rangifer tarandus* | GCA_019903745.1 |
| Siberian musk deer | *Moschus moschiferus* | GCA_004024705.2 |
| Lesser kudu | *Tragelaphus imberbis* | GCA_006410775.1 |

For the Suidae outgroup, we included the following taxa:

| | Species name | NCBI Genome Accession |
|---|---|---|
| Chacoan peccary | *Catagonus wagneri* | GCA_004024745.2 |
| Camel | *Camelus dromedarius* | GCA_000803125.3 |
| Bottlenose dolphin | *Tursiops truncatus* | GCA_011762595.1 |

Each outgroup genome assembly was first split into small 500 bp chunks using faSplit[30], which were then aligned to the cattle/pig reference genome using `bwa mem` with default parameters. For each reference genome (cattle or pig) we merged resulting individual BAM files using samtools merge. This resulted in a single BAM for each reference genome containing aligned fragments from different outgroup species. A consensus fasta was then called for each BAM separately using the `-doFasta 2` option in ANGSD[31] which assigns the most common nucleotide across the outgroups. The state at each monomorphic site i.e. coded as A, T, G or C in the fasta file was used as ancestral state downstreams analyses.

**Calculating genomic load.** Genomic load was obtained by summing the impact of deleterious alleles across the genome of each individual. These analyses were restricted to distant relative alignment (based on cattle and pig reference genome) as the conservation scores needed to measure the level of deleteriousness were not available for the close relative reference genome (buffalo and babirusa). Genomes with average depth of coverage > 5x were included in this analysis following Benjelloun *et al.*[32].

Starting with the filtered SNP panels generated using graphtyper[12] (see population structure section above), we removed outgroups and low coverage individuals from each VCF and annotated ancestral alleles using VcfAncestralAllele from JVarKit[33]. We retained biallelic sites where the reference allele matched the ancestral allele using `bcftools view -i 'INFO/AA==REF' -v snps`[11]. Finally, to allow cross species comparison, only sites within genes that were 1:1 orthologous between cattle and pig (following Ensembl release 107) were included. In total 453,134 sites for anoa and 299,929 sites for babirusa were used for this analysis.

For each individual, we obtained genotype likelihoods for all possible genotype at all sites that met the above criteria directly from BAM files using ANGSD[31] with command options '-GL 2 -doGlf 4 -doCounts 1 -minQ 30 -minMapQ 30 -remove_bads 1'. Genotype probabilities were then calculated from the likelihoods employing a Bayesian framework with an informed prior as described in Plassais *et al*.[34].

As a quantitative measure for deleteriousness, we relied on three types of conservation score as proxies: SIFT[35], phastCons[36], and PhyloP[37] scores. SIFT scores were obtained from Ensembl based on the Variant Effect Predictor (VEP)[38] while phyloP and phastCons were obtained from UCSC Genome Browser[30] using liftOver to convert coordinates from the human genome build (hg38) to that of the cattle (Btau9) and pig (Sscr11) reference genome. Sites were considered deleterious if their score were above (or below depending on the type of score) a specific threshold: i.e. SIFT scores <0.05, phastCons scores ≥0.75, and phyloP scores ≥1.5. The 0.05 cutoff for SIFT was obtained following Ng and Henikoff[39], the phyloP 1.5 cutoff was obtained from Librado *et al*.[40]. The phastCons cutoff reflects the 5% upper tail of the joint distribution of binned scores across both reference genomes.

To calculate genomic load, we sum conservation score across sites weighted by the genotype probability of the segregating non-ancestral allele (see equation 4 and 6 below). In the case of phyloP and phastCons, which have only one score for any non-ancestral state, we simply multiplied the score with the genotype probability of the derived allele. In the case of SIFT scores, which are available for each possible allele at a site, we considered only the score corresponding to the derived allele. Given that SIFT scores represent a normalised probability that an amino acid change is tolerated, their values are negatively correlated with deleteriousness, thus, we rescaled the conservation score at site *i* for a allele *j* as:

$$C_i = 1 - SIFT_{ij} \tag{3}$$

Where $SIFT_{ij}$ is the SIFT score at site *i* for the derived allele *j*. This allowed us to compare more easily the load measures derived from the three scores (the higher the value, the stronger the impact of deleterious mutations on that individual).

In this study we considered two types of genomic load: the homozygous load ($L_{HO}$) where deleterious mutations are regarded as recessive and the total load ($L_T$) which represents the worst-case scenario in which all deleterious mutations are dominant. The homozygous load was calculated as follows:

$$L_{HO} = \frac{S_{HO}}{n} \tag{4}$$

$$S_{HO} = \sum_{i=1}^{n} \frac{C_i P_i(DD_i)}{P_i(DD_i) + P_i(AA_i) + P_i(AD_i)} = \sum_{i=1}^{n} C_i P_i(DD_i)$$

where *n* is the number of sites considered, $C_i$ is the conservation score at site *i*, $P_i(DD_i)$ is the the posterior probability of observing the homozygous derived genotype at site *i*, $P_i(AA)$ is the posterior probability of observing the homozygous ancestral genotype at site *i*, and $P_i(AD)$ is the posterior probability of observing the heterozygous genotype at site *i*.

Following the same logic we can calculate the heterozygous load ($L_{HE}$) as shown in equation (5):

$$L_{HE} = \frac{S_{HE}}{n} \tag{5}$$

$$S_{HE} = \sum_{i=1}^{n} \frac{C_i P_i(AD_i)}{P_i(DD_i)+P_i(AA_i) + P_i(AD_i)} = \sum_{i=1}^{n} C_i P_i(AD_i)$$

while the total load ($L_T$) is the sum of the homozygous and heterozygous load, i.e.:

$$L_T = \frac{S_{HE} + S_{HO}}{n} = \frac{\sum_{i=1}^{n} C_i P_i(DD_i) + C_i P_i(AD_i)}{n} \quad (6)$$

All calculations were performed using a custom python script which is available on https://github.com/sagitaninta/Wallacea/tree/main/Individual_Mutational_Load.

We used one-way ANOVAs, for each species (aov() in R version 4.3.3[18], to assess whether load varied significantly across regions using P value threshold of 0.005[41]. We found that total load varied significantly across regions, using all three scores (p<0.005; Table S2) except for anoa/phastCons (P = 0.017) and anoa/phyloP (P = 0.006) where the difference across populations is suggestive. Homozygous load scores varied significantly (p<0.005) across regions in all cases except for anoa/phastCons (P = 0.006). In both species, and using all three conservation scores, our results suggest that total load is higher in the larger population on mainland Sulawesi while homozygous load score is higher in small island populations (Togian/Buton; Figure 3; Figure S4).

**Constructing a site frequency spectrum (SFS) for deleterious positions.** We constructed a SFS for different classes of deleterious alleles to better characterise the degree to which purifying selection acted in each population. To do so, we used the Ensembl VEP[38] impact ratings, i.e., “low”, “modifier”, “moderate”, and “high”, which is indicative of the deleteriousness of a mutation e.g. synonymous are “low” while loss-of-function mutations are deemed to have a “high” impact[41].

We ran VEP on the VCF used for our load analysis, adding allele frequency (AF) using `bcftools +fill-tags` tags filtering out sites that were non-variable (i.e. fixed differences with outgroup). We then extracted the coordinates of sites with low, modifier, moderate, and high impact and kept only sites in one-to-one ortholog between cattle and pig. These sites were used as input to calculate a SFS for each impact category using ANGSD. We first estimated site allele frequency (SAF) likelihood (-C 50 -minQ 30 -minMapQ 30 -dosaf 1 -GL 2) for each of three populations of anoa and babirusa: North Sulawesi, Southeast Sulawesi, and Buton/Togean.

The number of sites included in the SFS analysis for anoa:

| | High | Moderate | Modifier | Low |
|---|---|---|---|---|
| North Sulawesi | 1,404 | 112,034 | 62,815 | 162,221 |
| Southeast Sulawesi | 1,388 | 110,293 | 61,784 | 160,516 |
| Buton | 1,392 | 111,024 | 62,137 | 161,096 |

and babirusa:

| | High | Moderate | Modifier | Low |
|---|---|---|---|---|
| North Sulawesi | 811 | 71,032 | 47,195 | 118,455 |
| Southeast Sulawesi | 791 | 69,924 | 46,532 | 117,348 |
| Togean | 777 | 69,447 | 46,151 | 116,689 |

Ancestral alleles were specified during this step to allow polarisation of alleles and obtain an unfolded SFS. We then used ANGSD’s realSFS function to obtain the maximum likelihood estimate of the SFS.

**Composite likelihood test for purifying selection.** We compared the shape of the SFS for mutations of different categories (low, modifier, moderate, high, etc.) to assess the strength of purifying selection in different populations. If the population experienced strong purifying selection, the SFS of the higher impact mutation will shift from the expectation i.e. the distribution expected under neutrality. More specifically, we used a composite likelihood (CL) approach to compare two SFS, similar to Nielsen *et al.*[43]. In Nielsen *et al*.[43], the composite likelihood (CL) value of a SFS can be calculated by multiplying the probability of observing a derived allele at frequency $\boldsymbol{j}$ where $\boldsymbol{1} \leq \boldsymbol{j} \leq \boldsymbol{n-1}$, denoted as $\boldsymbol{p_j}$, along $\boldsymbol{n}$ chromosomes in a population as:

$$CL(p) = \prod_{j=1}^{n-1} {p_j}^{k_j}$$

where $k_j$ is the number of SNPs with derived allele at frequency $j$. The maximum likelihood estimates of $p_j$ is $\frac{k_j}{k}$.

Here we compared SFS of the low impact category (assumed to be neutral) to that of each other categories (modifier, moderate, high) as:

$$log(CL(p)) = \sum_{j=1}^{n} k_j \cdot log(p_j)$$

where $k_j$ is then the count of the derived alleles at frequency $j$ at sites in the modifier, moderate, high impact classes, and the maximum likelihood of $p_j$ is the proportion of low impact sites at frequency $j$. This composite likelihood was computed separately for each impact class.

We employed a bootstrapping approach (1000 replicates) to estimate uncertainty of the observed difference between the CL values of the SFS of the four impact ratings. To address the varying number of sites across impact categories, we resampled (with replacement) 1000 sites from each category to match the the lowest number of sites found in a population/impact category (i.e. high impact). Then, to increase the clarity in the visualisation, we normalised the CL values by dividing them with the mean of the 1000 bootstrap values of the low impact CL to show the relative magnitude of shift of the SFS from the three other impact ratings (modifier, moderate, high) compared to the expected SFS (low).

**Methods - Habitat Suitability modelling**

**Environmental predictor covariates.** We first evaluated the 19 bioclimatic variables (averaged over 1970-2000) and associated elevation data (derived from the SRTM data) from WorldClim v2.1[44]. We downloaded these variables for the geographic limits of Sulawesi (and surrounding small islands - Buton and Togean) using the "raster" package in R[45]. All environmental layers were clipped and masked to the species-specific area of interest: Sulawesi plus Buton for anoa, and Sulawesi plus the Togean islands for babirusa. Temperature variables were then processed as standard by dividing by a factor of 10 and elevation data was converted to a terrain ruggedness index (TRI) using the "spatialEco" R package[46]. This resulted in a set of 20 variables which were tested for collinearity using the variance inflation factor (VIF) to remove the highly correlated variables (th=3) with the "usdm" R package[47].

**Forest cover.** We used a forest cover layer downloaded from the Global Forest Change repository (v1.6; Hansen *et al.*[48]) and processed following procedures in Voigt *et al*.[49] to represent the primary and secondary forest categories used by the Indonesian Ministry of Forestry in the year 2018). Briefly, forest is defined as stands >5 ha with a natural composition and structure that had not been cleared in recent history (before the year 2000) and having >70% tree canopy cover at the Landsat pixel (30 m resolution) scale. This layer was clipped and masked to the species-specific area of interest and resampled to match the extent of the environmental predictors using the nearest neighbour method in the R "raster" package[45].

**Occurrences records.** For both anoa and babirusa, presence-only occurrence localities were compiled using all possible records from museums, previous studies (Frantz *et al.*[3]) and GBIF[50-51], a total of 263 occurrences for anoa and 514 occurrences for babirusa. These records were then filtered to those with accurate and reliable coordinates and retaining only one unique record per coordinate to reduce bias from clumping. Next, we constrained occurrences to the forest cover layer as both taxa are forest dependent. This resulted in a final set of 70 records each of anoa and babirusa (Data S1 & additional records in Data S3). We extracted the predictor covariates retained in the final model, at these coordinates, with the extract function in the "raster" package[44].

**Habitat suitability modelling.** We generated an ensemble species distribution model (ESDM) for each taxon based on the filtered environmental variables and the occurrence records using the "SSDM" package in R[52]. We tested eight algorithms: Artificial neural network (ANN), Classification tree analysis (CTA), Generalised additive model (GAM), Generalised boosted regression models (GBM), Generalised linear model (GLM), Multivariate adaptive regression splines (MARS), Random Forest (RF) and Support vector machines (SVM). Each algorithm was repeated 10 times and evaluated using the AUC metric with a threshold of 0.70 (Table S3). We also extracted the environmental variables that were retained in the model and their importance in determining the model. We penalised the ESDM projections post-hoc by forest cover to account for recent changes from deforestation. This was achieved by converting non-forested pixels, from the ESDM, to zero values, and resulted in the final habitat suitability scores constrained by forest cover for both taxa.

**Categorical habitat class.** The forest cover constrained ESDM model was then reclassified based on the percentiles of habitat suitability scores (e.g. 20% quantile intervals). In this way the model was divided into five habitat classes: from least suitable habitat (the lowest 20% of habitat suitability values) to most suitable habitat (top 20% of habitat suitability values) - class 1 = 0-20%, class 2 = 20-40%, class 3 = 40-60%, class 4 = 60-80%, class 5 = 80-100%. For each class, the corresponding patches were extracted using the 'landscape metrics' package in R[53] and converted to binary rasters (based on the maximum and minimum values). Using QGIS version 3.34.2[54], all layers were projected to the Asia South Albers Equal Area coordinate reference system (ESRI:102028). Each of the 5 categorical layers per taxa, were clipped to regional polygons corresponding to the north, southeast, and east Sulawesi, and the islands of Buton and Togean (Figure 2D). Although not a separate population cluster, the east has been kept as a separate region from the north, because of the distance and geography between these two peninsulas. The total area of each habitat class within these regions was calculated (Table S4) using the field calculator in QGIS[54].

**Representation of suitable habitat in various land uses.** Using the overlap analysis tool in QGIS, the proportion of each categorical habitat suitability class layer (1-5) that overlapped with different land-use categories was calculated (Figure S4). To do this we used a spatial layer where the land use across the region of interest had been assigned.
Unprotected land in Indonesia is broadly categorised for:

- (1) conversion (i.e. converted to non-forest land uses such as agriculture),
- (2) production (i.e. timber extraction, forestry), or protection.
- Areas under protection can be further classified as:
- (3) strict conservation areas (i.e. the highest levels of protection for biodiversity conservation as national parks and wildlife reserves, IUCN protected areas category I-II),
- (4) less strict conservation areas (i.e. protected for biodiversity conservation, but with little enforcement, IUCN categories III-V), or
- (5) watershed protection forest (i.e. forest outside the official conservation area network but set aside for watershed protection)[55].

Specifically, for Togean, we included the marine protected area (MPA) of Kepulauan Togean, which is included in the WDPA database[56] and in recent protected areas planning research[57]. This is because these islands represent important habitat for the babirusa, and the MPA extends onto the terrestrial land mass of Togean. In fact, it covers about 45.3% of the land (calculated in QGIS using the intersect and field calculator tools). This MPA is designated as an IUCN category II park and so the area covering terrestrial land was added to the strict PA category for the overlap calculations. Any other land classes on Togean, e.g. some small patches of production were removed, as this information is assumed to be no longer up to date.

**Land use and high-quality habitat.** By overlapping habitat suitability outputs with the land-use map we find that 25% of the most suitable anoa habitat (ie. the top quantile) and 16% of the most suitable babirusa habitat falls within conservation areas. Between the habitat classes, we see the largest amount of protected areas within the highest quality (Figure S4).

We used Chi-squared tests to compare the number of cells of each habitat class, which overlapped with either conservation areas or other land-uses. When comparing strictly IUCN protected with unprotected areas, we find a strong relationship with habitat class for anoa ($X^2$= 93.9, df = 4, p<0.05). Whereas for babirusa there was no significant relationship between number of cells and level of protection ($X^2$=8.0721, df = 4, p=0.088). However, when Hutan Lindung was included in the proportion of protected area (IUCN + Hutan Lindung) we found a significant relationship between the number of cells under protection compared to unprotected for both taxa (anoa: $X^2$= 41.38, df = 4, p<0.05, babirusa: $X^2$=144.68, df = 4, p<0.05). For both taxa, the Hutan Lindung significantly increases the amount of protection of the least suitable habitat (class 1, 0-20%) and not for the highest quality habitat classes.

**Supplementary figures**

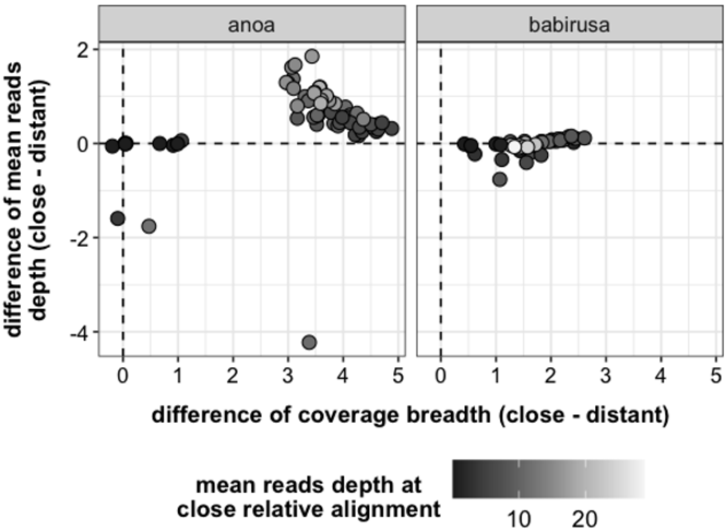


**Figure S1. Difference in mean reads depth and breadth of coverage using different genomes.** Close relative: *Bubalus bubalis* for anoa, *Babyrousa celebensis* for babirusa and distant relative: *Bos taurus* for anoa, *Sus scrofa* for babirusa.

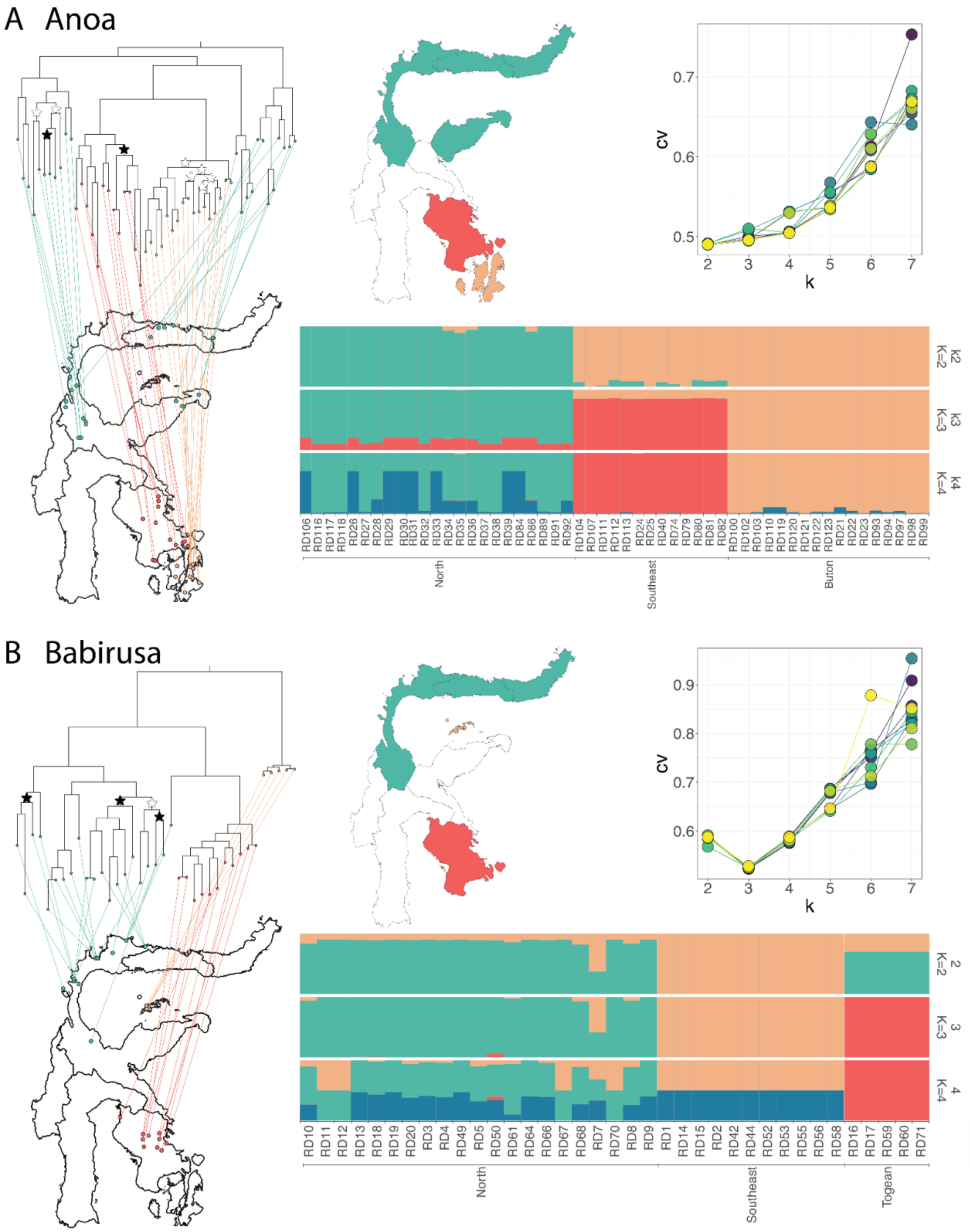


**Figure S2. Population structure analyses including maximum likelihood phylogenetic trees based on SNPs in genomes with >5x coverage and admixture analyses. (A)** anoa analyses based on 1,053,534 SNPs and 53 individuals, with phylogenetics tree indicating geographic location of sample (left), map of Sulawesi showing the colour for each geographic region (top, middle), admixture cross-validation error (top, right) and admixture plot for K2-4, where each unique ancestry is represented by a different coloured cluster. **(B)** babirusa analyses based on 1,011,533 SNPs and 37 individuals, with phylogenetic tree indicating geographic location of sample (left), map of Sulawesi showing the colour for each geographic region (top, middle), admixture cross-validation error (top, right) and admixture plot for K2-4, where each unique ancestry is represented by a different coloured cluster. Trees (A & B left) are rooted but the outgroups have been removed to aid plotting on the map. Most branches have very high support in both tests (UF bootstrap and SH-like ratio test >95) except those indicated with stars. Branches with white stars had high SH-like ratio test (>95) but low UF bootstrap support (70-78) while branches with black stars had low support in both tests (<74).

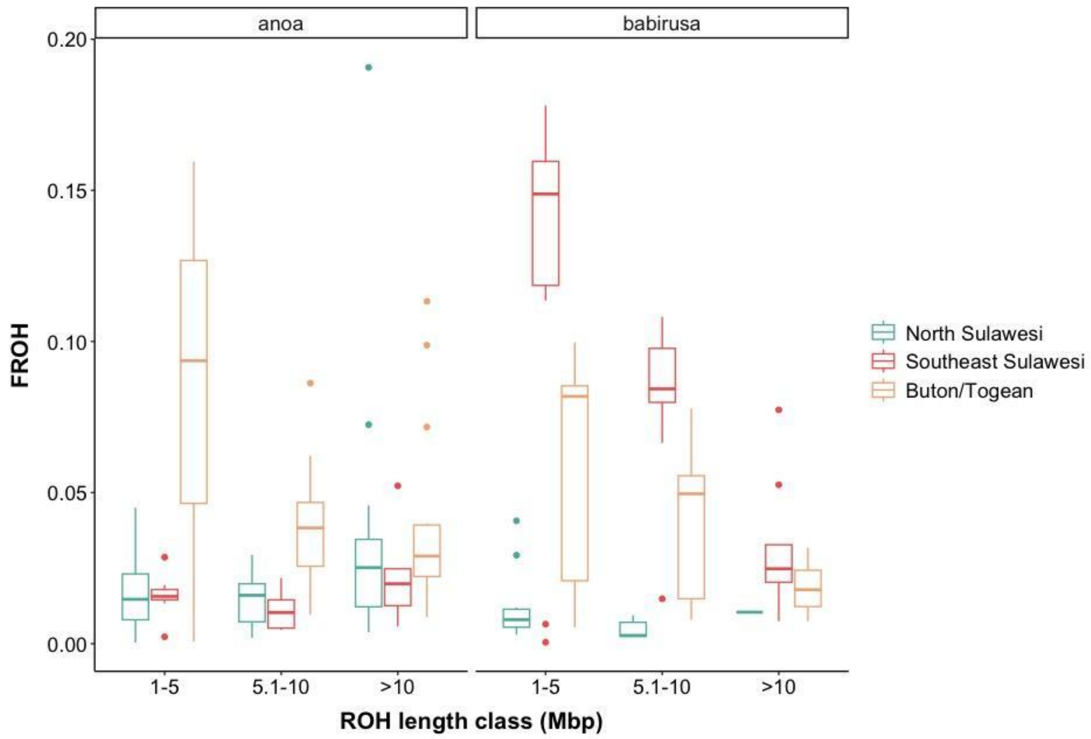


**Figure S3. Genomic inbreeding coefficient (FROH) at different ROH segment length groups of the different populations of anoa and babirusa.**

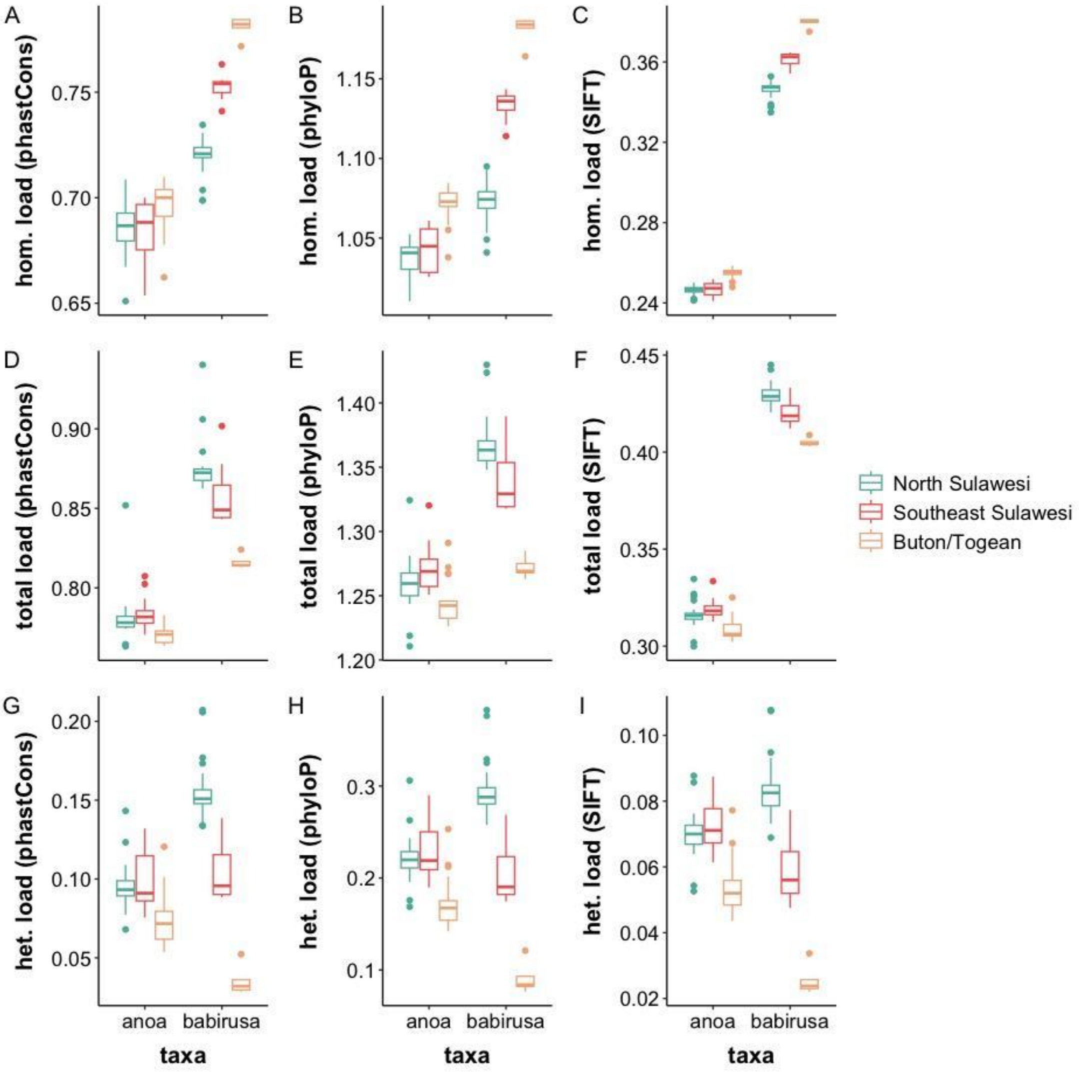


**Figure S4. Analysis of deleterious mutation load.** For anoa and babirusa calculated using phastCons**,** phyloP**,** and SIFT scores in **(A-C)** only homozygous, **(D-F)** both heterozygous and homozygous genotypes, and **(G-I)** only heterozygous sites.

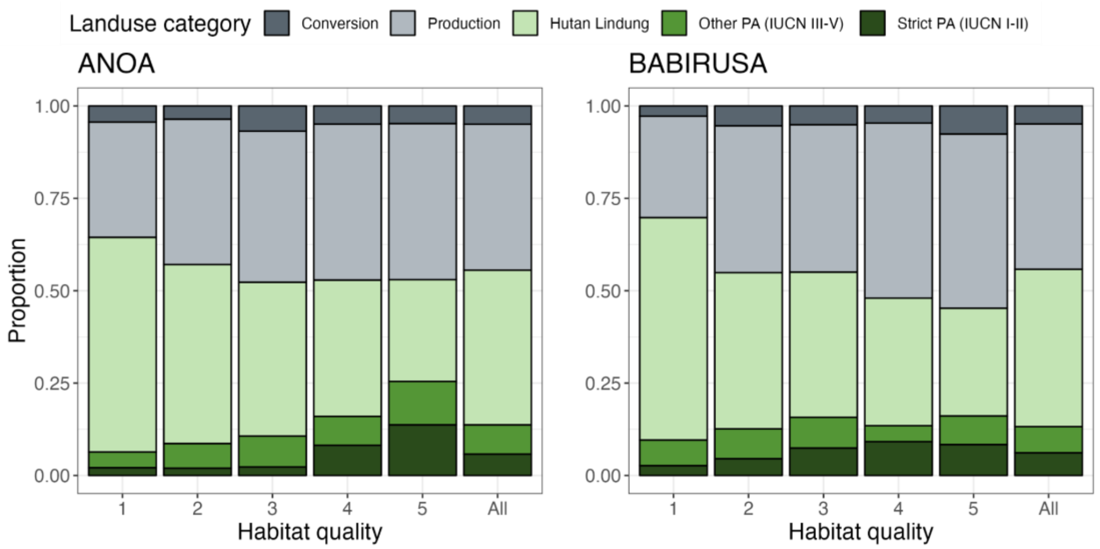


**Figure S5. Habitat suitability and land use category overlap.** Shows the proportion of the habitat suitability model that falls within areas of different land use categories for anoa (left) and babirusa (right). The models are split into quantiles based on the suitability scores where 1 = least suitable (0-20% of the habitat scores) and 5 = most suitable habitats (>80% of the habitat scores). The land-use categories are either non-protected: 1) conversion = converted land, 2) production = production forests associated with timber extraction and forestry, or protected to some degree 3) Hutan Lindung = forest protected for watersheds, not for biodiversity, 4) Other PA = IUCN category protected areas III-V, 5) Strict PA = IUCN protected areas category I-II.

**Supplementary Tables**

**Table S1. Results of one-way ANOVA testing the effect of ROH segment class length and population origin on the FROH per segment class of the anoa and babirusa populations.** P-values with asterisk mark significant effect of the variables in the model compared to null model (FROH ~ 1) using a threshold of 0.005 and the pairs with significant difference detected using post hoc TukeyHSD (P < 0.005) were shown in their respective tests (NO=North Sulawesi, SE=Southeast Sulawesi, BT=Buton, TO=Togean).

| **Model** | **df** | **F** | **P** | **Significantly different pairs (P<0.005)** |
|---|---|---|---|---|
| **Anoa** | | | | |
| **area** | 2 | 20.73 | * | BT-NO, BT-SE |
| **segment length class** | 2 | 3.2236 | 0.048 | N/A |
| **segment length class*area** | 8 | 10.079 | * | |
| area | 2 | 24.8853 | * | BT-NO, BT-SE |
| segment length class | 2 | 4.1500 | 0.018 | N/A |
| area:segment length class | 4 | 5.6394 | * | BT:1-5-NO:1-5,<br>BT:1-5-SE:1-5,<br>NO:5.1-10-BT:1-5,<br>SE:5.1-10-BT:1-5,<br>BT:5.1-10-BT:1-5,<br>NOi:>10-BT:1-5,<br>SE:>10-BT:1-5,<br>BT:>10-BT:1-5 |
| **Babirusa** | | | | |
| **area** | 2 | 19.266 | * | SE-NO |
| **segment length class** | 2 | 1.455 | 0.241 | N/A |
| **segment length class*area** | 8 | 12.211 | * | |
| area | 2 | 29.453 | * | SE-NO, TO-SE |
| segment length class | 2 | 15.134 | * | (>10)-(1-5) |
| area:segment length class | 4 | 2.129 | 0.089 | SE:1-5-NO:1-5,<br>SE:5.1-10-NO:1-5,<br>NO:5.1-10-SE:1-5,<br>TO:5.1-10-SE:1-5<br>SE:>10-SE:1-5<br>TO:>10-SE:1-5,<br>SE:5.1-10-NO:5.1-10 |

**Table S2**. **Results of one-way ANOVA testing the effect of population origin on the abundance of deleterious sites of different states in the different conservation scores in anoa and babirusa.** P-values with asterisk mark significant effect of region of origin using a threshold of 0.005 and the pairs with significant difference detected using post hoc TukeyHSD (P < 0.005) were shown in their respective tests (NO=North Sulawesi, SE=Southeast Sulawesi, BT=Buton, TO=Togean).

| **Score** | **Taxa** | **Sites** | **F** | **df** | **P-value** | **Significantly different pairs** |
|---|---|---|---|---|---|---|
| phastCons | anoa | homozygous | 4.75 | 2 | 0.013 | N/A |
| | | total | 4.79 | 2 | 0.013 | N/A |
| | babirusa | homozygous | 144.40 | 2 | * | NO-SE, NO-TO, SE-TO |
| | | total | 25.86 | 2 | * | NO-TO, SE-TO |
| phyloP | anoa | homozygous | 41.63 | 2 | * | NO-BT, SE-BT |
| | | total | 7.03 | 2 | 0.002 | SE-BT |
| | babirusa | homozygous | 203.80 | 2 | * | NO-SE, NO-TO, SE-TO |
| | | total | 39.42 | 2 | * | NO-SE, NO-TO, SE-TO |
| SIFT | anoa | homozygous | 48.89 | 2 | * | NO-BT, SE-BT |
| | | total | 9.66 | 2 | * | SE-BT |
| | babirusa | homozygous | 156.70 | 2 | * | NO-SE, NO-TO, SE-TO |
| | | total | 38.14 | 2 | * | NO-SE, NO-TO, SE-TO |

**Table S3. Ensemble Species Distribution Model (ESDM) evaluation.** Algorithms maintained in the final ensemble model for each of the taxa based on the area under the curve metric (AUC) at a threshold level of 0.70, including the uncorrelated variable importance, for those variables maintained in the taxa specific ensemble models (Pearson's correlation).

| | **Anoa** | **Babirusa** |
|---|---|---|
| **Model accuracy (AUC)** | | |
| **Full ensemble model** | **0.752<br>(7 algorithms, 7 covariates)** | **0.747<br>(4 algorithms, 7 covariates)** |
| GAM | 0.75 | - |
| GLM | 0.70 | - |
| SVM | 0.77 | 0.76 |
| MARS | 0.76 | - |
| GBM | 0.77 | 0.75 |
| CTA | 0.74 | 0.76 |
| RF | 0.76 | 0.71 |
| **Variable importance of those retained in the final model:** | | |
| BIO2 - Mean diurnal range | 17.9 | - |
| BIO3 - Isothermality | - | 8.2 |
| BIO7 - Temperature annual range | 7.5 | 21.1 |
| BIO8 - Mean temperature of wettest quarter | 5.4 | - |
| BIO9 - Mean temperature of driest quarter | - | 14.2 |
| BIO13 - Precipitation of wettest month | 31.0 | - |
| BIO14 - Precipitation of driest month | 18.8 | - |
| BIO15 - Precipitation seasonality | - | 23.6 |
| BIO16 - precipitation of the warmest month | 8.4 | - |
| BIO18 - Precipitation of warmest quarter | - | 10.2 |
| BIO19 - Precipitation of coldest quarter | - | 9.8 |
| TRI | 11.0 | 12.9 |

**Table S4. Proportion of each class of the model (class 1 - 5) that falls within the geographic regions for each taxa.** The classes are based on the habitat quantile of the model with class 1 being the least suitable (lowest 20% quantile) and class 5 being the most suitable (top 20% quantile). The regions north, southeast and east Sulawesi based on the map and zones of endemism from Frantz et al[3] (Figure 1).

| **Habitat suitability** | **Anoa** | | | | **Babirusa** | | |
|---|---|---|---|---|---|---|---|
| | **North Sulawesi** | **Southeast Sulawesi** | **East Sulawesi** | **Island - Buton** | **North Sulawesi** | **Southeast Sulawesi** | **Island - Togean** |
| Unclassified - Unforested | 0.21 | 0.21 | 0.12 | 0.44 | 0.22 | 0.23 | 0.40 |
| Class 1 - very poor (0-20%) | 0.02 | 0.11 | 0.04 | 0.00 | 0.03 | 0.08 | 0.00 |
| Class 2 - poor (20-40%) | 0.05 | 0.27 | 0.18 | 0.002 | 0.09 | 0.16 | 0.05 |
| Class 3 - moderate (40%-60%) | 0.11 | 0.24 | 0.38 | 0.04 | 0.19 | 0.16 | 0.09 |
| Class 4 - good (60-80%) | 0.30 | 0.09 | 0.15 | 0.07 | 0.21 | 0.19 | 0.05 |
| Class 5 - very good (80-100%) | 0.31 | 0.08 | 0.13 | 0.45 | 0.26 | 0.18 | 0.41 |

### Dataset legends

**Dataset S1 (separate file).** Provenance of anoa and babirusa samples used in this study including the region and coordinates origin.

**Dataset S2 (separate file).** Publicly available bovid and suid genomes used in this study.

**Dataset S3 (separate file).** Additional records of occurrence of anoa and babirusa used to generate the ESDM.
